# Effects of arm weight support on neuromuscular activation during reaching in chronic stroke patients

**DOI:** 10.1101/668046

**Authors:** Keith D Runnalls, Pablo Ortega-Auriol, Angus J C McMorland, Greg Anson, Winston D Byblow

## Abstract

To better understand how arm weight support (WS) can be used to alleviate upper limb impairment after stroke, we investigated the effects of WS on muscle activity, muscle synergy expression, and corticomotor excitability (CME) in 13 chronic stroke patients and 6 age-similar healthy controls. For patients, lesion location and corticospinal tract integrity were assessed using magnetic resonance imaging. Upper limb impairment was assessed using the Fugl-Meyer upper extremity assessment with patients categorised as either mild or moderate-severe. Three levels of WS were examined: low=0, medium=50 and high=100 % of full support. Surface EMG was recorded from 8 upper limb muscles, and muscle synergies were decomposed using non-negative matrix factorisation from data obtained during reaching movements to an array of 14 targets using the paretic or dominant arm. Interactions between impairment level and WS were found for the number of targets hit, and EMG measures. Overall, greater WS resulted in lower EMG levels, although the degree of modulation between WS levels was less for patients with moderate-severe compared to mild impairment. Healthy controls expressed more synergies than patients with moderate-severe impairment. Healthy controls and patients with mild impairment showed more synergies with high compared to low weight support. Transcranial magnetic stimulation was used to elicit motor-evoked potentials (MEPs) to which stimulus-response curves were fitted as a measure of corticomotor excitability (CME). The effect of WS on CME varied between muscles and across impairment level. These preliminary findings demonstrate that WS has direct and indirect effects on muscle activity, synergies, and CME and warrants further study in order to reduce upper limb impairment after stroke.

## Introduction

Stroke is a leading cause of acquired adult disability with two-thirds of stroke survivors experiencing lingering upper limb impairment (Feigin et al., 2010; Mendis, 2013). The likelihood of regaining functional independence after stroke is strongly influenced by the initial severity of motor deficits and subsequent recovery of motor function (Kwakkel et al., 1996; Patel et al., 2000; Meijer et al., 2003; Veerbeek et al., 2011). Conventional therapy attempts to engage mechanisms of motor learning to reshape control of the remaining neuromechanical repertoire. Task-specific recovery of upper limb function is facilitated when physical therapy exercises are performed with a high number of repetitions (Kwakkel et al., 2004; Veerbeek et al., 2014). However, high repetition schedules are not always achieved (Lang et al., 2009). Providing arm weight support (WS) may augment the performance of arm movements and increase the dose of therapeutic exercise that is possible (Kwakkel and Meskers, 2014). Studies of WS as an adjuvant to neurorehabilitation have typically included WS as a component of robotic-aided therapies without separating it from other assistive or resistive forces and sensory feedback (Johnson, 2006; Loureiro et al., 2011). Less is known about the separable effects of WS on the upper limb movements of stroke patients (Prange et al., 2009a; Krabben et al., 2011). A better understanding of WS and its underlying neural mechanisms may inform the application of WS in stroke rehabilitation.

In addition to facilitating greater training dosages, WS can also improve movement quality. During reaching tasks, WS reduces antagonist muscle activity in both healthy older adults and chronic stroke patients (Prange et al., 2009a; 2009b). Abnormal coupling of joint torques between the shoulder and elbow is also lessened with WS (Dewald and Beer, 2001; Beer et al., 2004). The stereotyped flexor synergy can thus be mitigated with WS, permitting greater elbow extension and access to the reaching workspace (Beer et el., 2007; Sukal et al., 2007). Taken together, it appears that WS may facilitate a dissociation of strength and motor control deficits. Understanding transient modulation of motor control with WS has relevance because the expressed patterns of neuromotor activity may be reinforced with repetition.

Muscle synergies identified through decomposition of recorded EMG can provide insight into the underlying structure of neuromotor activity. Differences in the recruitment or activation of synergies might reflect context-specific or compensatory motor control, whereas differences in synergy structure might reflect more enduring neuroanatomical constraints. Data obtained from stroke patients performing a dynamic upper limb task confirm that the internal structure of synergies can be preserved despite altered movement performance (Cheung et al., 2009). In contrast, a change in synergy structure was observed in an isometric task where the three heads of the deltoid muscle were consistently expressed as a single synergy and the extent of its activation was related to the degree of impairment (Cheung et al., 2012). Patients with more impairment exhibited fewer synergies, reflecting a lower dimensional neuromechanical repertoire. In healthy adults the level of WS influenced the activation, but not composition, of muscle synergies during a reaching task (Coscia et al., 2014). However, the effect of WS on muscle synergies following stroke, and its interaction with impairment, has not been adequately investigated.

Corticomotor excitability (CME) across the upper limb is modulated with the amount of WS in healthy adults (Runnalls et al., 2014; 2015; 2017). The reduction of antigravity torques required for shoulder abduction has indirect effects on other upper limb muscles through putative neural linkages. To what extent muscle activity and CME are sensitive to partial WS in chronic stroke patients is unknown.

In the present study, we investigated the effects of WS in chronic stroke patients with a range of upper limb impairment. First, we examined upper limb muscle activations during a reaching task. Surface electromyography (EMG) was recorded from eight muscles while participants performed reaching movements to an array of targets with high, medium, and low levels of WS. We expected that greater WS would allow patients to reach more targets. It was hypothesised that support level would interact with impairment severity and target location to modulate integrated EMG area (iEMG) across the upper limb. Second, we conducted a muscle synergy analysis on the EMG data recorded during the reaching task. We hypothesised that patients with more severe impairment would express fewer synergies. We also expected that the application of WS would permit the expression of more synergies. Last, we examined CME at high, medium, and low levels of WS. Transcranial magnetic stimulation (TMS) was used to elicit motor evoked potentials (MEPs) during a static shoulder abduction task. CME was examined by comparing stimulus-response (SR) curves fitted to means derived from statistical models of MEP area. It was hypothesised that WS would modulate CME, and the pattern of modulation would depend on impairment severity.

## Methods

### Participants

Thirteen chronic stroke patients (mean age 70.8 years, range 47–88 years, four females) with upper limb impairment participated in this study (Table 1). Patients were included if they reported any degree of upper limb impairment resulting from a first-ever stroke that occurred more than six months before testing. Patients were excluded if they had no active range of motion at the shoulder. Patients were excluded from the MRI or TMS component if screening revealed any contraindications. Patients were characterised as having mild impairment if the upper extremity Fugl-Meyer assessment score was 50 or more. Patients with scores below 50 were characterised as moderate-severe (mod-sev). Six neurologically healthy adults (mean age 65.2 years, range 51–71 years, all right dominant, two females) participated as age-similar controls. All participants gave written informed consent. Study procedures were approved by the University of Auckland Human Participants Ethics Committee in accordance with the Declaration of Helsinki.

**Table 1.** Participant characteristics

| Participant | Age (y) | Sex | Chronicity (months) | Affected Limb | FM | ARAT | MAS | FA<sub>AI</sub> | Lesion Location | Group |
| --- | --- | --- | --- | --- | --- | --- | --- | --- | --- | --- |
| 1 | 77 | M | 50 | Right | 66 | 57 | 0 | -0.06 | Left Cerebral White Matter | Mild |
| 2 | 80 | M | 49 | Left | 45 | 49 | 1 | 0.13 | Right Cerebral White Matter | Mod-Sev |
| 3 | 53 | M | 40 | Right | 62 | 57 | 0 | NA | Left Cerebral White Matter | Mild |
| 4 | 70 | F | 90 | Right | 66 | 57 | 0 | 0.04 | Left Cerebral Cortex | Mild |
| 5 | 74 | M | 144 | Right | 12 | 0 | 3 | 0.08 | Left Cerebral White Matter | Mod-Sev |
| 6 | 72 | M | 104 | Right | 56 | 57 | 1.5 | 0.05 | Left Cerebral White Matter | Mild |
| 7 | 83 | F | 43 | Right | 65 | 57 | 0 | 0.08 | Left Cerebral Cortex | Mild |
| 8 | 72 | F | 84 | Left | 9 | 0 | 3 | 0.42 | Right Cerebral White Matter | Mod-Sev |
| 9 | 47 | M | 39 | Right | 40 | 37 | 2 | 0.06 | Left Cerebral White Matter | Mod-Sev |
| 10 | 71 | F | 27 | Left | 16 | 3 | 3 | 0.17 | Right Cerebral White Matter | Mod-Sev |
| 11 | 88 | M | 71 | Left | 58 | 45 | 0 | 0.16 | Right Cerebral White Matter | Mild |
| 12 | 74 | M | 13 | Right | 56 | 53 | 1 | 0.15 | Left Cerebral White Matter | Mild |
| 13 | 59 | M | 167 | Left | 17 | 3 | 1 | 0.31 | Right Cerebral White Matter | Mod-Sev |
| **Min** | 47 |  | 13 |  | 9 | 0 | 0 | -0.06 |  |  |
| **Max** | 88 |  | 167 |  | 66 | 57 | 3 | 0.42 |  |  |
| **Mean** | 70.8 | 9 M \| 4 F | 70.8 | 8 R \| 5 L | 43.7 | 36.5 | 1.2 | 0.13 | 2 Cortical \| 11 Subcortical | 7 Mild \| 6 Mod-Sev |
| 14 | 71 | M |  |  |  |  |  |  |  | Control |
| 15 | 69 | M |  |  |  |  |  |  |  | Control |
| 16 | 59 | F |  |  |  |  |  |  |  | Control |
| 17 | 51 | M |  |  |  |  |  |  |  | Control |
| 18 | 71 | M |  |  |  |  |  |  |  | Control |
| 19 | 70 | F |  |  |  |  |  |  |  | Control |
| **Min** | 51 |  |  |  |  |  |  |  |  |  |
| **Max** | 71 |  |  |  |  |  |  |  |  |  |
| **Mean** | 65.2 | 4 M \| 2 F |  |  |  |  |  |  |  | 6 Control |

**Note:** FM, upper extremity Fugl-Meyer score (maximum 66). ARAT, action research arm test (maximum 57). MAS, modified Ashworth spasticity scale for the elbow (maximum 4). FA<sub>AI</sub>, fractional anisotropy asymmetry index for the posterior limb of the internal capsule (perfect symmetry = 0). Mild, FM > 50; Moderate-Severe (Mod-Sev), FM < 50

### Magnetic resonance imaging

Lesion location and corticospinal tract integrity were assessed using magnetic resonance imaging (Figure 1). Brain images were acquired using a 3T MAGNETOM Skyra MRI scanner (Siemens, Germany). An MP-RAGE sequence was used to acquire high-resolution T_1_-weighted anatomical images (T_R_ = 1900 ms, T_E_ = 2.07 ms, FoV = 256 mm, voxel dimensions of 1.0×1.0×1.0 mm). Diffusion-weighted images (DWI) were acquired using a single shot echo planar imaging sequence (T_R_ = 3600 ms, T_E_ = 92.4 ms, FOV = 220 mm, voxel dimensions 2.0×2.0×2.0 mm), with thirty diffusion gradient orientations (b = 2000 s/mm^2^).

**Fig 1.**
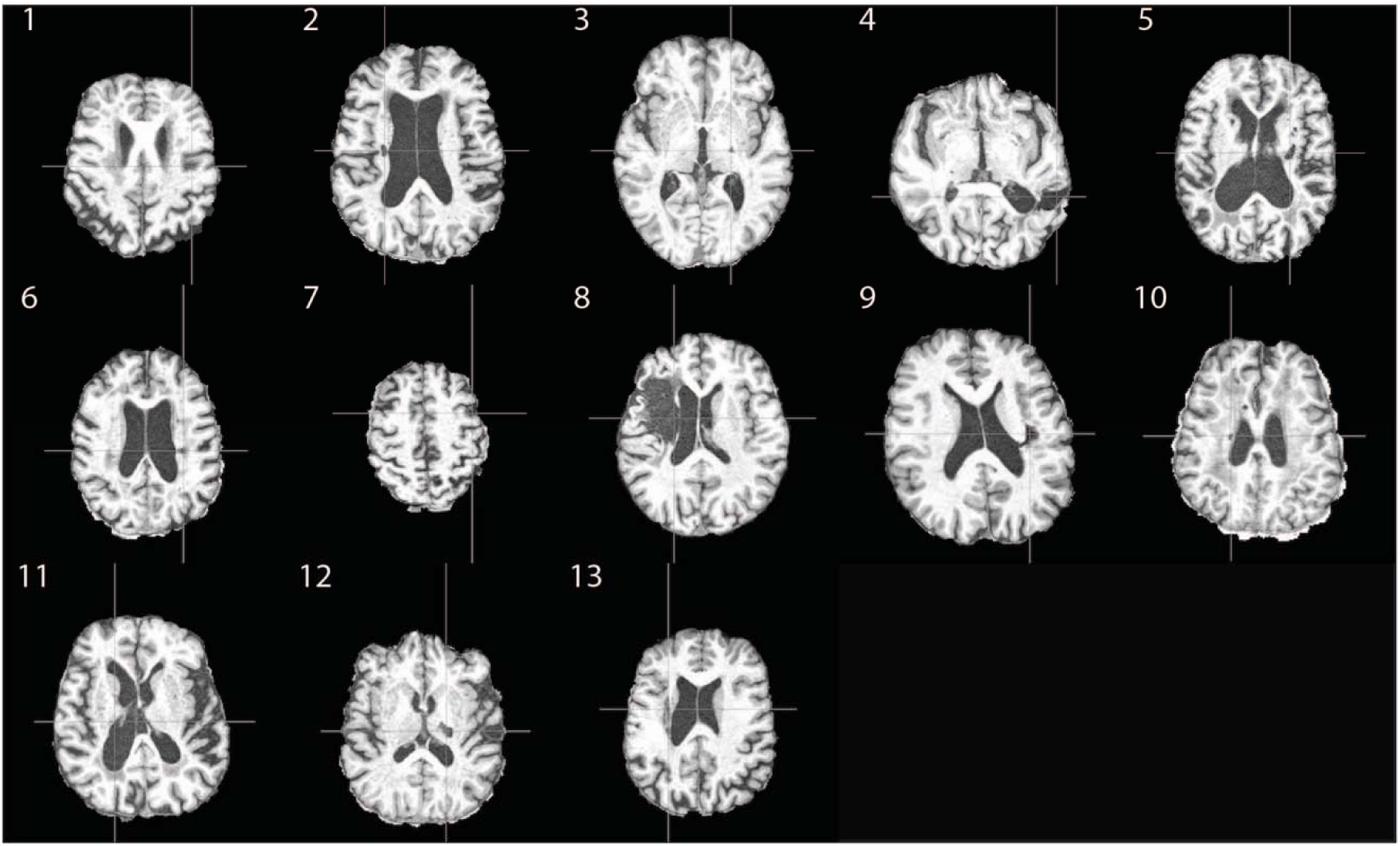
Anatomical T_1_-weighted images in the transverse plane at the level of the lesion for each patient. Patient numbers correspond with Table 1

Lesions were located and masked on T_1_-weighted images using FSLView from the FMRIB Software Library (Jenkinson et al., 2012). Diffusion-weighted images were processed using FMRIB’s Diffusion Toolbox. Images were skull stripped using the Brain Extraction Tool (Smith, 2002), and corrected for motion and eddy currents (Jenkinson and Smith, 2001; Jenkinson et al., 2002; Andersson and Sotiropoulos, 2016). To quantify corticospinal tract (CST) integrity, mean fractional anisotropy (FA) was calculated within the posterior limb of the internal capsule (PLIC) for the ipsilesional (FA_Ipsi_) and contralesional (FA_Contra_) hemispheres. A fractional anisotropy asymmetry index (FA_AI_) was calculated as FA_AI_ = (FA_Contra_ - FA_Ipsi_) / (FA_Contra_ + FA_Ipsi_), resulting in a value between −1 and 1 for each participant (Stinear et al., 2007). Positive values correspond to reduced ipsilesional FA.

### Session Order

Participants attended an initial session to complete clinical assessments with a physical therapist. The therapist was not involved in any other aspects of the study. In a subsequent session, participants performed the repeated measures reaching task. Four blocks of reaching trials were completed for high, medium, and low levels of arm weight support. The total twelve blocks were performed in a randomised order. Eligible participants then completed the TMS component in the same session. Single-pulse TMS was used to obtain stimulus-response curves at high, medium, and low levels of arm weight support while participants maintained a static arm posture. Six blocks of stimulation (two for each level of weight support) were performed in a quasi-randomised order. In a separate session, MRI was used to obtain anatomical and diffusion-weighted images (DWI) of the brain. Healthy control participants did not undergo clinical assessments or MRI.

### Posture and arm support

The reaching task and TMS measures were completed while participants sat in a chair with their feet on the floor and unsupported arm resting on their lap. Arm weight support was provided to the stroke-affected limb, or dominant limb for healthy controls, by a SaeboMAS arm support system (Saebo Inc., Charlotte, NC). Force was provided by spring tension through a brace that cradled the proximal forearm. The forearm was secured in the brace using elasticized fabric wrap. A standardised static arm position was set with the shoulder flexed forward approximately 80° and abducted 45° in the horizontal plane, the elbow flexed at 90°, and the forearm pronated palm down. In this position, the hand was in front of the shoulder with the elbow pointing laterally. The brace prevented rotation in the vertical plane ensuring the forearm was parallel to the floor. Joint angles were initially set using a goniometer and subsequently maintained by aligning the hand to a reference object.

Three levels of arm weight support were defined relative to the force required to compensate for the weight of the arm completely. At low support (0%), the device carried its weight but provided no additional support to the arm. Individualised levels of support were determined using a force titration procedure. Participants maintained the standardised static arm posture while supportive force was incrementally decreased from a magnitude that required voluntary shoulder adduction. High support (100%) was defined as the last point before root mean square EMG amplitude (rmsEMG) in the anterior deltoid was observed to deflect away from the baseline activity that persists even with excessive support (Runnalls et al. 2014; Runnalls et al. 2015; Runnalls et al. 2016). Medium support was then defined as 50% of high support.

### Electromyography

Surface electromyography data were recorded from eight muscles of the supported arm and hand: anterior deltoid (AD), middle deltoid (MD), posterior deltoid (PD), clavicular head of pectoralis major (PM), biceps brachii (BB), lateral head of triceps brachii (TB), brachioradialis (BRD), and extensor carpi radialis (ECR). Self-adhesive Ag-AgCl electrodes (Blue Sensor N; Ambu, Denmark) were placed approximately 2 cm apart in a bipolar montage over the belly of each muscle. The common ground electrode was placed over the acromion process (Red Dot; 3M Health Care, Canada). Signals were amplified (Grass P511AC; Grass Instrument Division, West Warwick, RI) with 1000x gain, band-pass filtered (10–1000 Hz), sampled at 2 kHz using a 16-bit A/D acquisition system (National Instruments, Austin, TX), and saved for offline analysis.

### Reaching task

Participants were seated facing a table-mounted robotic arm (UR5; Universal Robots, Denmark). A push-button assembly was mounted to the tool attachment point of the robot. The 6 cm diameter pushbutton responded to input anywhere on its surface; i.e. finger extension was not required. The robot moved the button to predefined locations to present it as the target for each reach. A trial would begin from a static start position in which the arm was adducted close to the torso with the elbow flexed at 90° and forearm oriented forward orthogonal to the coronal plane. If a participant could not reliably adopt this position, the nearest approximation was used.

Each block of trials was composed of the same sequence of fourteen targets (Figure 2). The targets were located at incrementally greater distances away from the start position along four direction vectors. The distribution was designed to probe the reachable limits of the forward workspace volume. Three targets were located along a low-wide vector, and four targets were located along a high-narrow vector. These vectors were mirrored laterally to test both lateral (ipsilateral) and medial (contralateral) directions.

**Fig 2.**
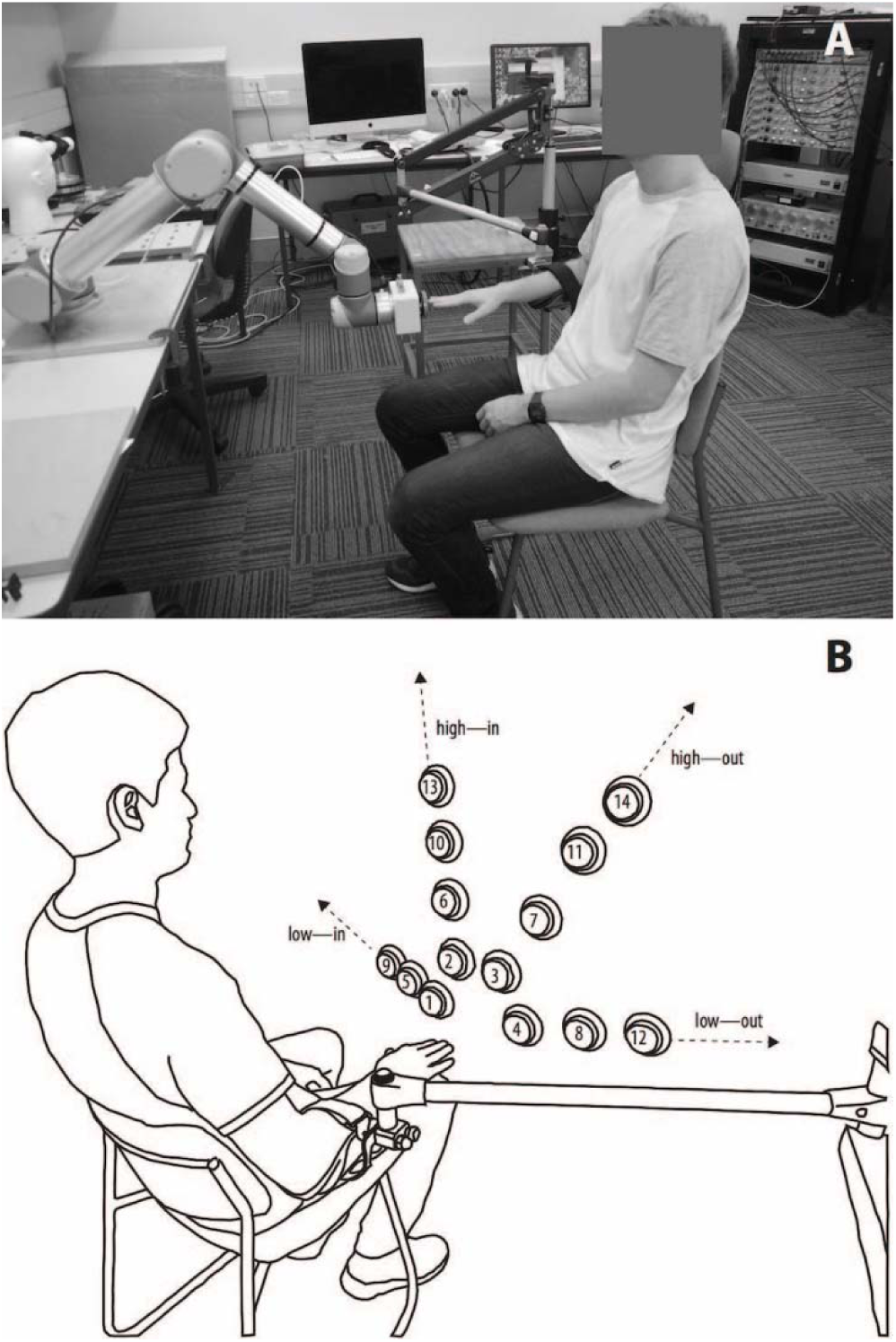
**A:** Demonstration of static start position with the robot presenting the push-button as a calibration point. **B:** Schematic illustration of reaching target positions. Targets were presented in numerical order. Targets were 10, 15, 20, and 25 cm anterior to the start position; 0, 10, 20, 30 and 40 cm above the start position; 5, 10, 15, 20, and 30 cm medial/lateral to the start position

A computer-generated tone simultaneously started data recording and cued the participant to begin the movement. Participants were instructed to keep their back against the chair and reach to push the button at a comfortable speed. Recording terminated when the button was pressed. A trial was flagged as incomplete if the target button could not be pressed without compensatory strategies such as forward torso lean or stabilisation with the unaffected arm.

Individual EMG traces were detrended, rectified, and low-pass filtered using a fourth-order zero-lag Butterworth filter with a cut-off frequency of 6 Hz using MATLAB R2013a (MathWorks, Natick, MA). The resulting EMG traces from individual muscles were inspected in parallel for each trial and trimmed as necessary to make the EMG onset time consistent between trials. Integrated EMG area (iEMG) was calculated as the dependent measure for each trace.

### Statistical Analyses

Statistical analyses of the reaching task were conducted using R 3.3.2 (R Core Team, 2016) with the *lme4*: Linear Mixed-Effects Models using ‘Eigen’ and S4 (Bates et al., 2015) and *car*: Companion to Applied Regression (Fox and Weisberg, 2010) packages. For each participant, iEMG was normalised between zero and one across conditions within each muscle. Linear mixed effects models were fitted for each muscle to investigate the effects of weight support, impairment severity, and target parameters on muscle activity. In each case, iEMG was modelled as the dependent variable with fixed effects for SUPPORT LEVEL, IMPAIRMENT, TARGET DISTANCE, TARGET HEIGHT, and TARGET SIDE. Random intercepts were included for SUBJECT. Model terms were tested using type II Wald F tests with Kenward-Roger degrees of freedom.

### Muscle synergy analysis

Raw EMG traces were detrended, rectified, and normalised to the maximum value across all muscles within each trial. Normalised traces were low-pass filtered using a fourth-order zero-lag Butterworth filter with a cut-off frequency of 6 Hz, then resampled to a length of 1000 points using shape-preserving piecewise cubic interpolation. For each combination of subject and support level, traces were averaged across repetitions to each target and concatenated to an N by T matrix D, where N is the number of EMG channels and T is the number of successfully reached targets multiplied by 1000 (number of data points of EMG trace). D was used as an input to the Non-negative Matrix Factorization Algorithm (MATLAB 2016b). NMF was modelled as D = W × C, where D is the original data set, W is the synergies, and C is the activation coefficient. The algorithm converged onto a constrained solution from 20 consecutive iterations with a difference between iterations of less than 0.01% of the calculated ‖D – W × C‖.

The NMF procedure takes as an input parameter the pre-defined number of modules or synergies to extract, between 1 and N. Dimensionality reduction of the original matrix is accomplished by selecting a minimum number of synergies that can reconstruct the matrix D to a certain threshold of quality, quantified by the variance accounted for (VAF). VAF is defined in Equation 1, where ODS is the variance of the original data set and RD is the variance of the reconstructed data. The number of synergies was determined when VAF > 90% to reconstruct D, and VAF > 80% to reconstruct each EMG channel.

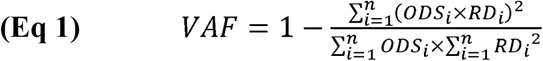

A mixed effects ANOVA (3 IMPAIRMENT x 3 SUPPORT LEVEL) was performed on the number of synergies. Mauchly’s test of sphericity was performed and degrees-of-freedom were adjusted using a Greenhouse-Geisser correction (□ = 0.62).

### Synergy Clustering

To compare synergy structures, it is necessary to associate equivalent synergies for each support level and impairment group. Normalized synergies from all participants within a SUPPORT LEVEL and IMPAIRMENT GROUP were pooled. Cluster analysis was applied to pooled synergies using K-medoids (Park and Jun, 2009), using the scalar product as the distance metric between clusters, and Silhouette index (Kaufman and Rousseeuw, 1990) to determine the correct number of clusters. For each resultant cluster a mean synergy was calculated, mean synergies of each IMPAIRMENT GROUP across SUPPORT LEVEL were then pooled together. To match similar synergies across SUPPORT LEVEL, a second cluster analysis was applied to the mean pooled synergies.

### Synergy comparison

To determine if synergies were conserved through different support levels, the scalar product was calculated between normalized synergies. The scalar product is a measure of similarity between vectors ranging from 0 (completely different) to 1 (identical). The positivity constraint of NMF means that even two synergies extracted from random data will have non-zero similarity values. Two synergies were defined as similar when their scalar product was above a threshold of the 95th percentile of a z-distribution of scalar products generated by comparing shuffled synergies (Ortega-Auriol et al, 2018). We calculated the distribution of similarity values using synergies constructed from pooled weight values that were randomly shuffled across epochs and muscles.

### Transcranial magnetic stimulation

Single-pulse TMS was delivered to M1 using a MagStim 200 magnetic stimulator (Magstim, Dyfed, UK). A figure-of-eight coil (Magstim D70^2^) was held tangentially to the scalp and angled to direct current in a posterior to anterior direction across the central sulcus. The coil was positioned at the optimal site for eliciting MEPs in the contralateral BB and ECR muscles and the location was marked on the scalp. Task motor threshold (MT) was defined as the minimum stimulus intensity that elicited a 100 µV MEP in four out of eight trials with the arm in the standardised static position at the high support level. All TMS was conducted while participants performed the static arm abduction task, i.e. actively maintaining the standardised static arm position.

Stimulus–response (SR) curves were collected for high, medium, and low levels of weight support. A single stimulation site was used to concurrently elicit MEPs in all muscles. Five stimulus intensities were set relative to task motor threshold of BB: −5, +5, +15, +25, and +35 % of maximum stimulator output (% MSO). For each curve, forty stimuli were delivered over two blocks (eight stimuli for each of the five intensities). To mitigate fatigue, participants rested their arm after every four stimuli as required.

Raw EMG traces were inspected and processed using Signal 5.11 (CED, Cambridge, UK). Trials were excluded from further analysis if there was no stimulus artefact or if there was phasic muscle activity present. Dependent measures were obtained from individual EMG traces. MEP area was calculated over a 20 ms window determined manually for each muscle. Background muscle activity was calculated as the rmsEMG amplitude over a 50 ms window preceding the stimulus.

Statistical analyses of background muscle activity and MEP area were conducted using R 3.3.2 with the lme4, car, and predictmeans: Calculate Predicted Means for Linear Models packages (Luo et al., 2014). Outliers were identified for each muscle by analysing background muscle activity on a within-participant basis. Observations of rmsEMG more than 1.5× the interquartile range either above the third quartile or below the first quartile, along with their associated MEP values, were removed from the dataset. MEP area was normalised between zero and one for each participant and muscle, across conditions. Logarithmic transforms were applied to normalised MEP area within the models to better satisfy the assumption of normally distributed residuals.

To assess the effect of WS on background muscle activity, as well as any interaction with impairment severity, linear mixed effects models were fitted for each muscle. In each case, BACKGROUND MUSCLE ACTIVITY was modelled as the dependent variable with fixed effects for SUPPORT LEVEL and IMPAIRMENT. Random intercepts were included for SUBJECT. Model terms were tested using type II Wald F tests with Kenward-Roger degrees of freedom.

For MEP area, independent linear mixed effects models were constructed for each muscle. In each case, MEP area was modelled as the dependent variable with fixed effects for STIMULUS INTENSITY, SUPPORT LEVEL, and IMPAIRMENT. BACKGROUND MUSCLE ACTIVITY (rmsEMG) was included as a continuous covariate term. The error term included random slopes for BACKGROUND MUSCLE ACTIVITY and random intercepts for SUBJECT. The models were then used to predict means and standard errors for MEP area at the median value of the background muscle activity distribution (Welham et al., 2004). This procedure controlled for systematic differences in background muscle activity thus permitting unbiased analysis of MEP area.

Stimulus-response curves were fitted to group level predicted means using nonlinear regression in Prism 7 (GraphPad, San Diego, CA). For each combination of support level and impairment, a three parameter Boltzmann function was fitted to predicted mean MEP area as a function of relative stimulus intensity (Devanne et al., 1997). The upper plateau was constrained to its theoretical range between zero and one to improve the rate at which regression converged on a fit. The slope and half-maximal stimulus intensity (S50) parameters were unconstrained. To test whether changes in support level shifted the stimulus-response curve, extra sum-of-squares F tests were used to assess whether individual regression curves fit the data significantly better than a single curve within each impairment group.

## Results

### Reaching task

Data from all nineteen participants (six healthy control, seven mild impairment, six moderate-severe impairment) were included in the analysis. The number of targets hit are presented in Figure 3. Control participants successfully hit all 14 targets at all support levels. Participants with mild upper limb impairment hit an average of 12.8 targets at low support and 12.9 at medium and high support. Those with moderate-severe upper limb impairment hit an average of 8.9 targets at low support, 10.6 at medium, and 11.4 at high. As expected, ANOVA indicated an interaction between IMPAIRMENT and SUPPORT LEVEL (*F*_(4,32)_ = 8.63, *p* = 0.002). Means for iEMG are presented in Figure 4 for each target. Muscles were analysed separately with independent statistical models. Targets were grouped by distance, side, and height to test contrasts. Supplementary Table 1 presents the results of the ANOVA for each muscle.

**Fig 3.**
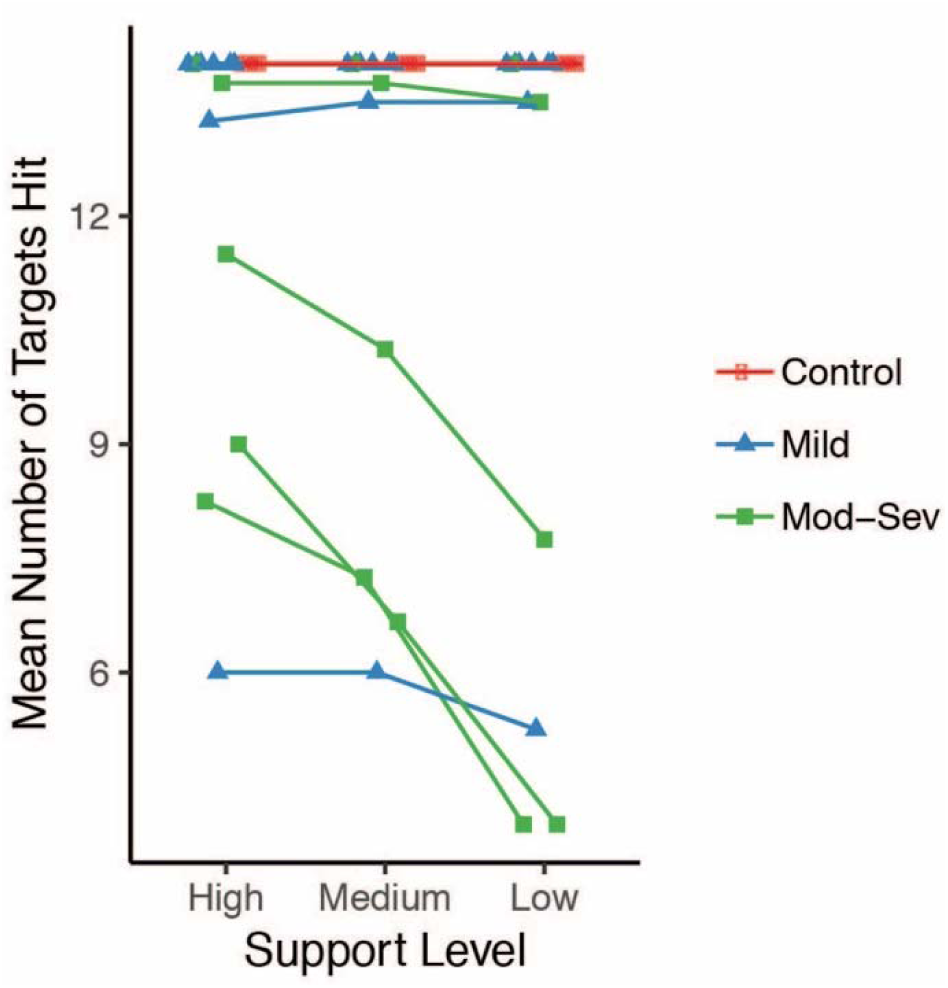
Mean number of targets hit at each support level for each participant. Colours represent impairment groups. Support level is a discrete variable and data points have been dodged horizontally for visualization only. Error bars indicate ±1 SEM

**Fig 4.**
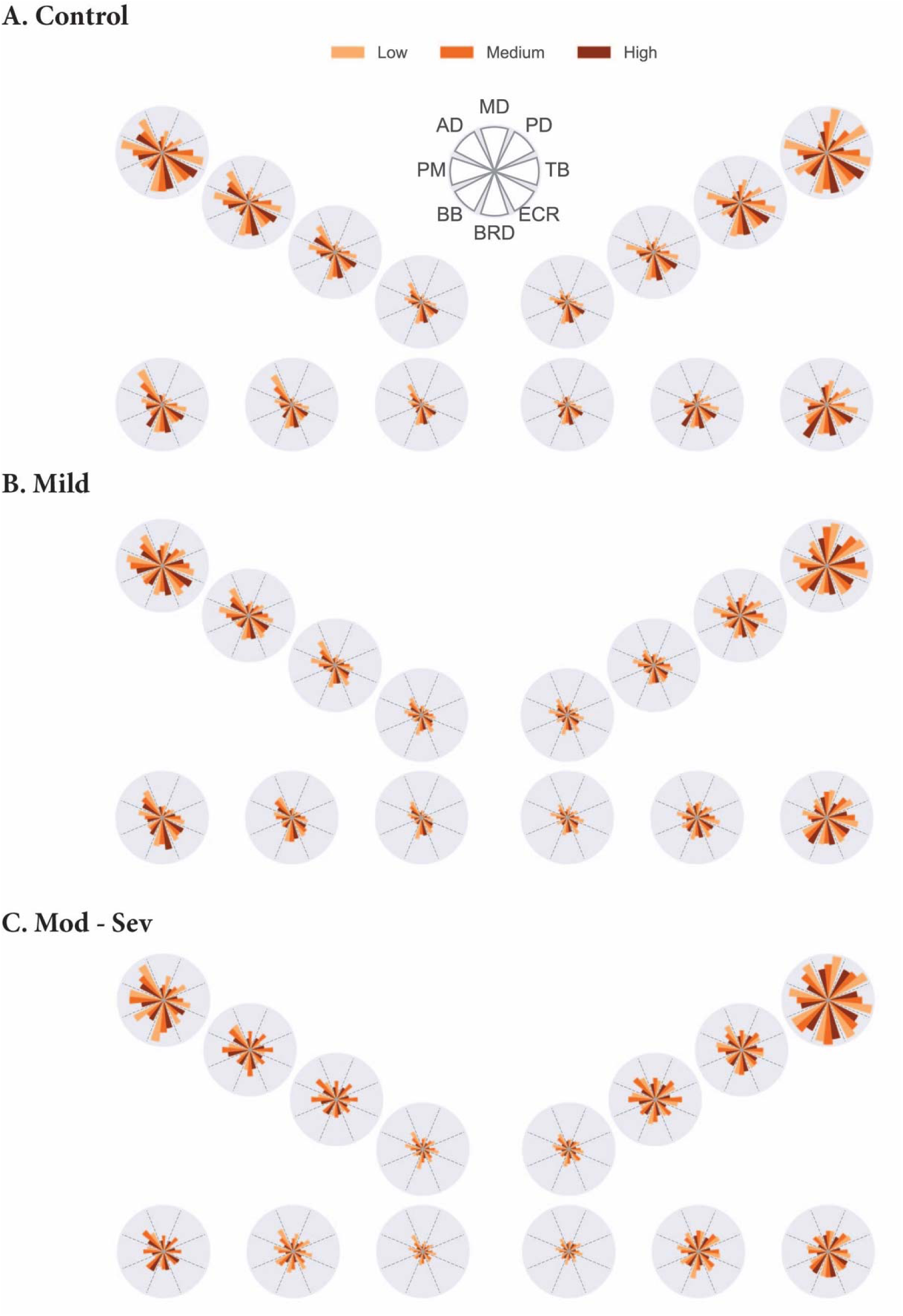
Mean normalised iEMG for **A:** Control, **B:** Mild, and **C:** Moderate-Severe impairment groups. Each circular subplot corresponds to a reaching target as presented in Figure 2B. Muscles are represented as sectors of each circle. Support level is indicated by colour

### Effect of weight support on number of synergies

The number of synergies identified for each IMPAIRMENT and SUPPORT LEVEL are presented in Figure 5. There was no interaction between IMPAIRMENT and SUPPORT LEVEL for the number of synergies, but there were main effects of both factors (Table 2). Planned contrasts revealed a greater number of synergies with high support (2.2, SD 1.0) compared to low support (1.6, SD 0.6; *F*_(1, 16)_ = 7.0, *p* = 0.018), with no difference between high and medium support levels (1.8, SD 0.6; *F*_(1, 16)_= 1.61, *p* = 0.22). The control group expressed more synergies (2.4, SD 0.7) compared to patients with moderate-severe upper limb impairment (1.3, SD 0.8; *d* = 1.111, *p* = 0.02) while there was no difference between mild (1.86, SD 0.4) and moderate-severe (1.3, SD 0.4; *d* = 0.52, *p* = 0.082) impairment levels. Given that patients were unable to reach all targets, subsets of the control group data were analysed for comparison to both the four and eight most common targets reached by the patient groups. A one-way repeated measures ANOVA conducted on the four target subset revealed no differences for the number of synergies observed in the control group across SUPPORT LEVEL (*F*_(1.1, 5.4)_ = 1.86, *p* = 0.21, □ = 0.55).

**Table 2.**
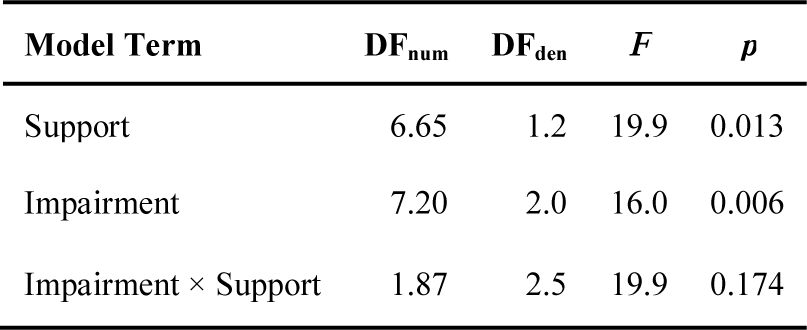
ANOVA for number of synergies expressed in reaching task

**Fig 5.**
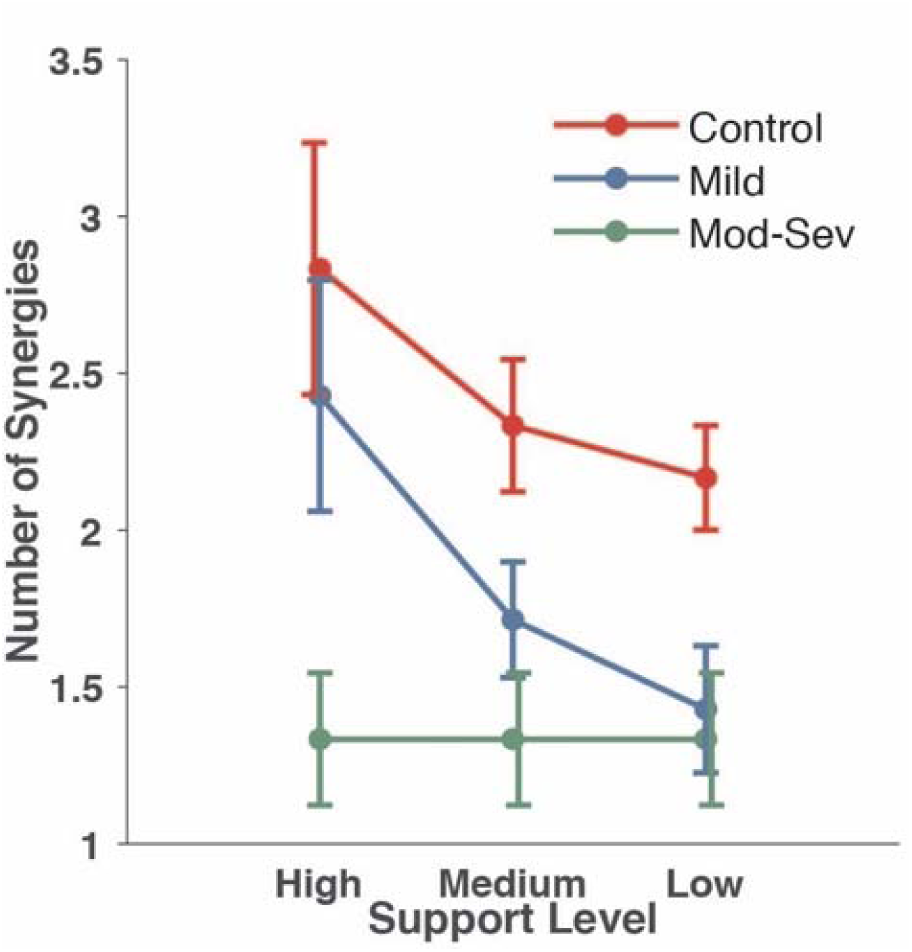
Mean number of extracted synergies at each support level for each impairment group. Colours represent impairment groups. Error bars indicate ±1 SEM

### Functional significance of synergies

The functional significance of a single synergy relates to the muscles with the highest weight in its structure. Three synergies were identified across the different groups and support levels. Although not all muscles were measured, the functional interpretation considered target positions and low weight of some muscles within a synergy structure: S1 or external rotation synergy, characterised by higher weights of AD and MD with an almost absent PM, S2 or internal rotation synergy with high weight of AD and PM, and S3 or flexion synergy with a higher weight of ECR, BR and BB.

### Synergy similarity

Synergy structures identified by the clustering analysis are presented in Figure 6. The similarity of synergies was quantified as the average pairwise scalar product within each cluster. The scalar product is a global measure of similarity across synergies. It is possible that small differences between synergies in vector space could reflect larger differences at the behavioural level. Thus, despite the similarities that we found across groups, clustered synergies that were different from the control group clusters were denoted as atypical.

**Fig 6.**
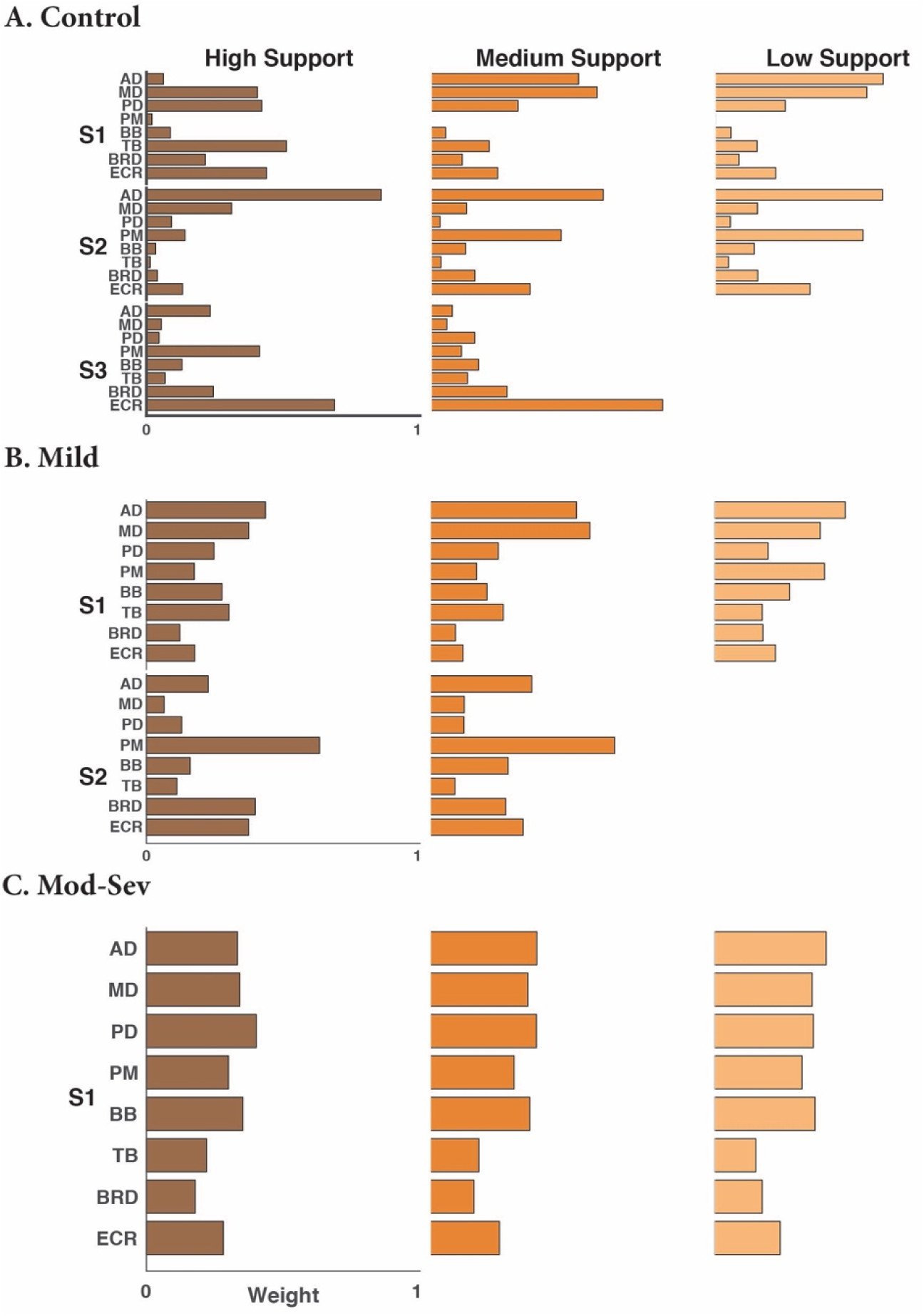
Structure of synergy clusters at different support levels. **A:** Controls, three synergies were able to reconstruct the data of the high and medium support, while only two were necessary for the low support level. Synergy S1 represents muscles involved in external rotation, S2, internal rotation, and S3, flexion. Synergy structure was conserved through the different support levels. **B:** Mild Impairment, two synergies were extracted under the high and medium support conditions, while only one synergy was identified with low support. **C:** Moderate-Severe Impairment, a single synergy was present across all three support levels

For the control group, cluster analysis revealed three clusters of similar synergies for high and medium support levels, and two for the low support level. The S1 synergy was present across all support levels with a moderate similarity (0.75, SD 0.1). The S2 (internal rotation) synergy was present across all support levels with a high mean similarity (0.81, SD 0.1). Finally, the S3 (flexion) synergy was only present for the high and medium support levels, and its mean similarity was moderate (0.78, SD 0.1). For patients with mild impairment there were two clusters across support levels: S1 (which appears functionally atypical compared to the control S1) was present at all support levels with a moderate similarity (0.71, SD 0.2). S2 (internal rotation) was present only for the high and medium support levels with high similarity (0.84, SD 0.2). Finally, the group with moderate-severe upper limb impairment presented a single (atypical) synergy for all support levels with a moderate similarity (0.77, SD 0.1). Between groups, synergy similarity was calculated on mean synergies across support levels. Similarities were high between S1-S1 (0.91, SD 0.1) and S2-S2 (0.85, SD 0.1) for the control and mild impairment groups respectively.

Mean pooled similarity across support levels for the mild impairment group was significantly higher than similarity across shuffled synergies (0.6, SD 0.1). A non-parametric ANOVA was used to determine differences between pooled synergy similarity of each group across support levels and the similarity of shuffled synergies. A Kruskal-Wallis test revealed the existence of differences across IMPAIRMENT (H_(3)_ = 172), and pairwise comparisons with Bonferroni-adjusted p-values found differences between the control, mild impairment, and moderate-severe impairment groups (all *p* < 0.001) and the shuffled synergies threshold.

### Transcranial magnetic stimulation

Data from six control, six mildly impaired, and two moderate-severely impaired participants were included in the analysis. Of the 40 stimuli delivered to each participant per condition, an average of 38.6 traces (range: 28–40) were retained in the final analysis. Traces were discarded based on outlying values of background muscle activity. Example EMG traces are presented in Figure 7.

**Fig 7.**
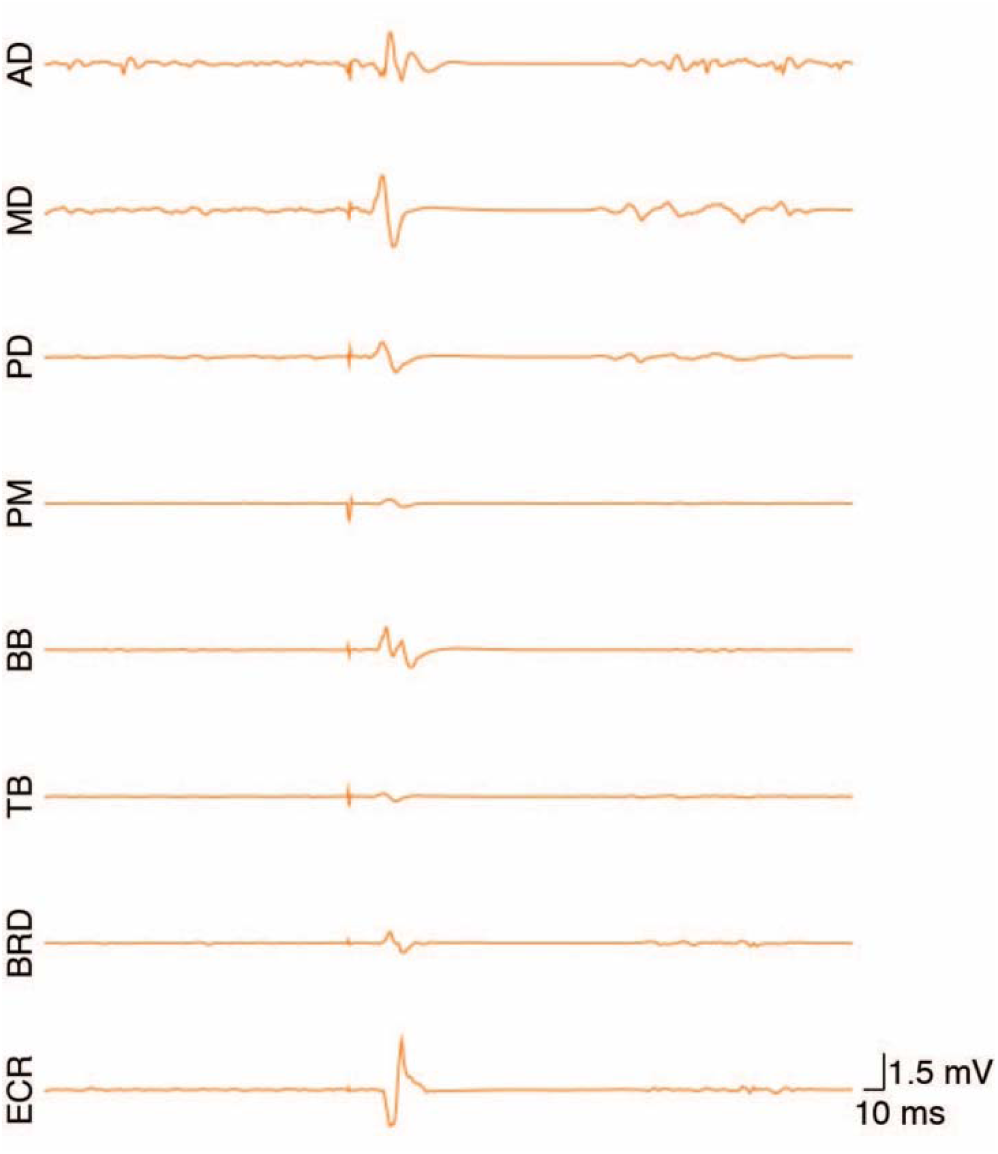
Single EMG traces showing motor evoked potentials from a representative participant using the medium support level. TMS intensity was set to task motor threshold + 25% MSO

### Effects of weight support and impairment on background muscle activity

Boxplots for background muscle activity are presented in Figure 8. All muscles exhibited a significant main effect of SUPPORT LEVEL and a significant interaction between SUPPORT LEVEL and IMPAIRMENT (Table 3). With the exception of BB, which exhibited less activity in the control group (0.008 mV) compared to mild (0.026 mV) and moderate-severe (0.021 mV) groups, there were no main effects of IMPAIRMENT.

**Table 3.** ANOVA for linear mixed models of background muscle activity in static abduction task

| **Muscle** | **Model Term** | **DF<sub>num</sub>** | **DF<sub>den</sub>** | **F** | **p** |
| --- | --- | --- | --- | --- | --- |
| **AD** | Support | 2 | 1620 | 907.30 | < .001 |
|  | Impairment | 2 | 11 | 2.30 | 0.147 |
|  | Support × Impairment | 4 | 1620 | 149.77 | < .001 |
| **MD** | Support | 2 | 1624 | 1140.23 | < .001 |
|  | Impairment | 2 | 11 | 0.12 | 0.892 |
|  | Support × Impairment | 4 | 1624 | 140.31 | < .001 |
| **PD** | Support | 2 | 1609 | 376.36 | < .001 |
|  | Impairment | 2 | 11 | 0.69 | 0.523 |
|  | Support × Impairment | 4 | 1609 | 141.06 | < .001 |
| **PM** | Support | 2 | 1589 | 370.35 | < .001 |
|  | Impairment | 2 | 11 | 2.41 | 0.135 |
|  | Support × Impairment | 4 | 1589 | 8.54 | < .001 |
| **BB** | Support | 2 | 1611 | 368.33 | < .001 |
|  | Impairment | 2 | 11 | 9.71 | 0.004 |
|  | Support × Impairment | 4 | 1611 | 36.16 | < .001 |
| **TB** | Support | 2 | 1597 | 242.27 | < .001 |
|  | Impairment | 2 | 11 | 3.09 | 0.086 |
|  | Support × Impairment | 4 | 1597 | 25.20 | < .001 |
| **BRD** | Support | 2 | 1581 | 89.47 | < .001 |
|  | Impairment | 2 | 11 | 2.42 | 0.135 |
|  | Support × Impairment | 4 | 1581 | 17.54 | < .001 |
| **ECR** | Support | 2 | 1588 | 19.00 | < .001 |
|  | Impairment | 2 | 11 | 2.90 | 0.097 |
|  | Support × Impairment | 4 | 1588 | 18.99 | < .001 |

**Fig 8.**
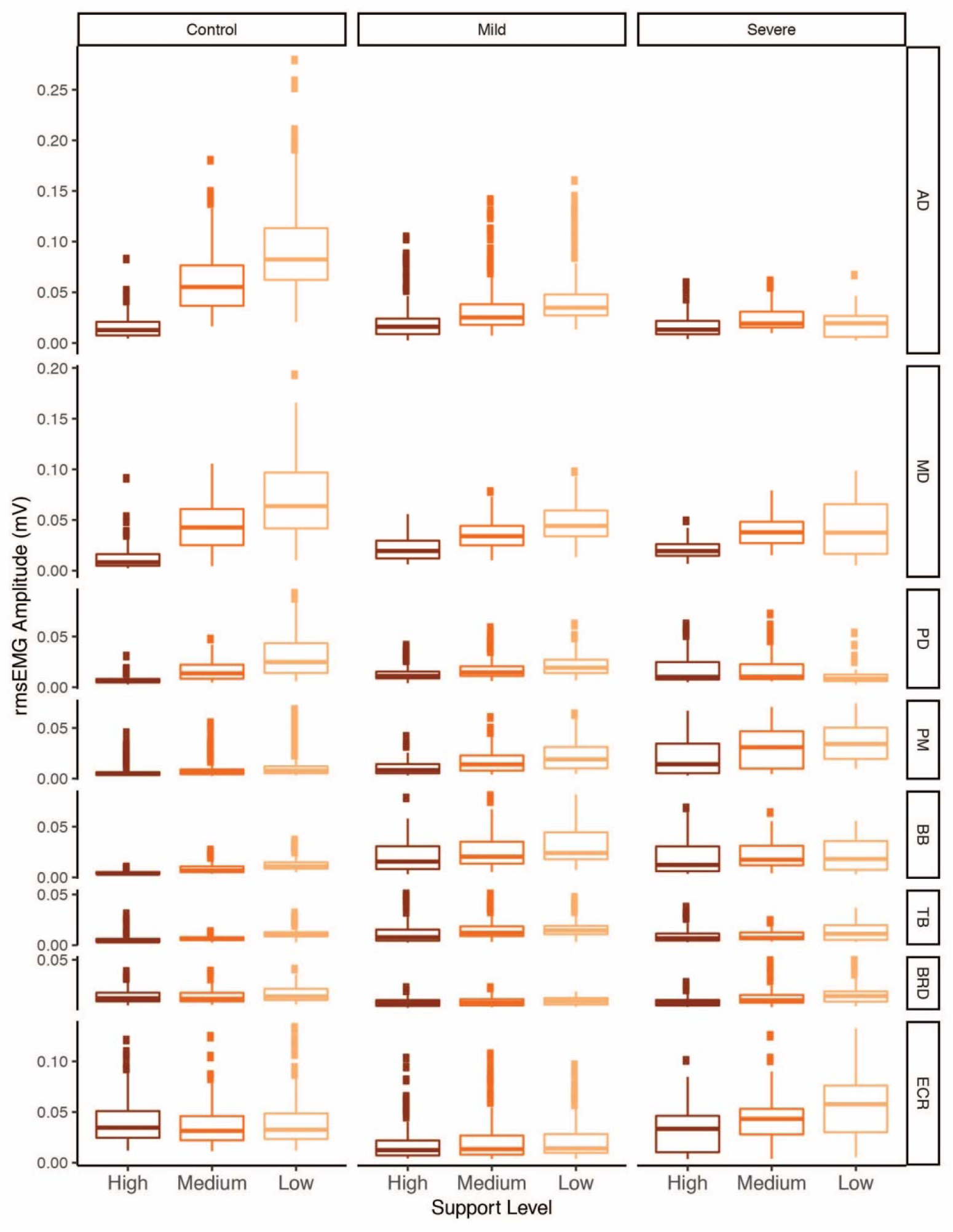
Background muscle activity at each support level for control, mild, and moderate-severe impairment groups during the standardised static arm abduction task. Boxplots summarise rmsEMG measured before each TMS stimulus

### Effects of weight support and impairment on MEP area and stimulus-response curves

Figure 9 presents stimulus-response data derived from the linear mixed effects models. Type II Wald χ^2^ tests of model terms indicated a significant effect (p < 0.001) of SUPPORT LEVEL on MEP area in all muscles. Mean normalized MEP area was then predicted for each combination of STIMULUS INTENSITY, SUPPORT LEVEL, and IMPAIRMENT. The procedure accounted for co-varying background muscle activity. Stimulus-response curves fitted to the predicted means were tested for differences between support levels. Results of the extra sum-of-squares F tests are presented in Table 4.

**Table 4.** Comparison of stimulus-response curve fits for support levels. Tests where p < 0.05 indicate different curves for each support level is the preferred model

| **Muscle** | **Impairment Group** | **$F_{(DFn,DFd)}$** | **$p$** |
| --- | --- | --- | --- |
| **AD** | Control | 100.10<sub>(6, 6)</sub> | 0.001 |
|  | Mild | 27.13<sub>(6, 6)</sub> | 0.001 |
|  | Mod-Sev | 0.99<sub>(6, 6)</sub> | 0.507 |
| **MD** | Control | 9.16<sub>(6, 6)</sub> | 0.008 |
|  | Mild | 19.88<sub>(6, 6)</sub> | 0.001 |
|  | Mod-Sev | 8.79<sub>(6, 6)</sub> | 0.009 |
| **PD** | Control | 61.08<sub>(6, 6)</sub> | <.001 |
|  | Mild | 7.68<sub>(6, 6)</sub> | 0.013 |
|  | Mod-Sev | 1.14<sub>(6, 6)</sub> | 0.438 |
| **PM** | Control | 8.26<sub>(6, 6)</sub> | 0.011 |
|  | Mild | 13.04<sub>(6, 6)</sub> | 0.003 |
|  | Mod-Sev | 19.62<sub>(6, 6)</sub> | 0.001 |
| **BB** | Control | 24.08<sub>(6, 6)</sub> | <.001 |
|  | Mild | 3.81<sub>(6, 6)</sub> | 0.064 |
|  | Mod-Sev | 1.01<sub>(6, 6)</sub> | 0.494 |
| **TB** | Control | 58.56<sub>(6, 6)</sub> | <.001 |
|  | Mild | 8.64<sub>(6, 6)</sub> | 0.009 |
|  | Mod-Sev | 1.14<sub>(6, 6)</sub> | 0.439 |
| **BRD** | Control | 9.81<sub>(6, 6)</sub> | 0.007 |
|  | Mild | 2.18<sub>(6, 6)</sub> | 0.183 |
|  | Mod-Sev | 5.68<sub>(6, 6)</sub> | 0.026 |
| **ECR** | Control | 1.78<sub>(6, 6)</sub> | 0.250 |
|  | Mild | 1.46<sub>(6, 6)</sub> | 0.329 |
|  | Mod-Sev | 36.66<sub>(6, 6)</sub> | <.001 |

**Fig 9.**
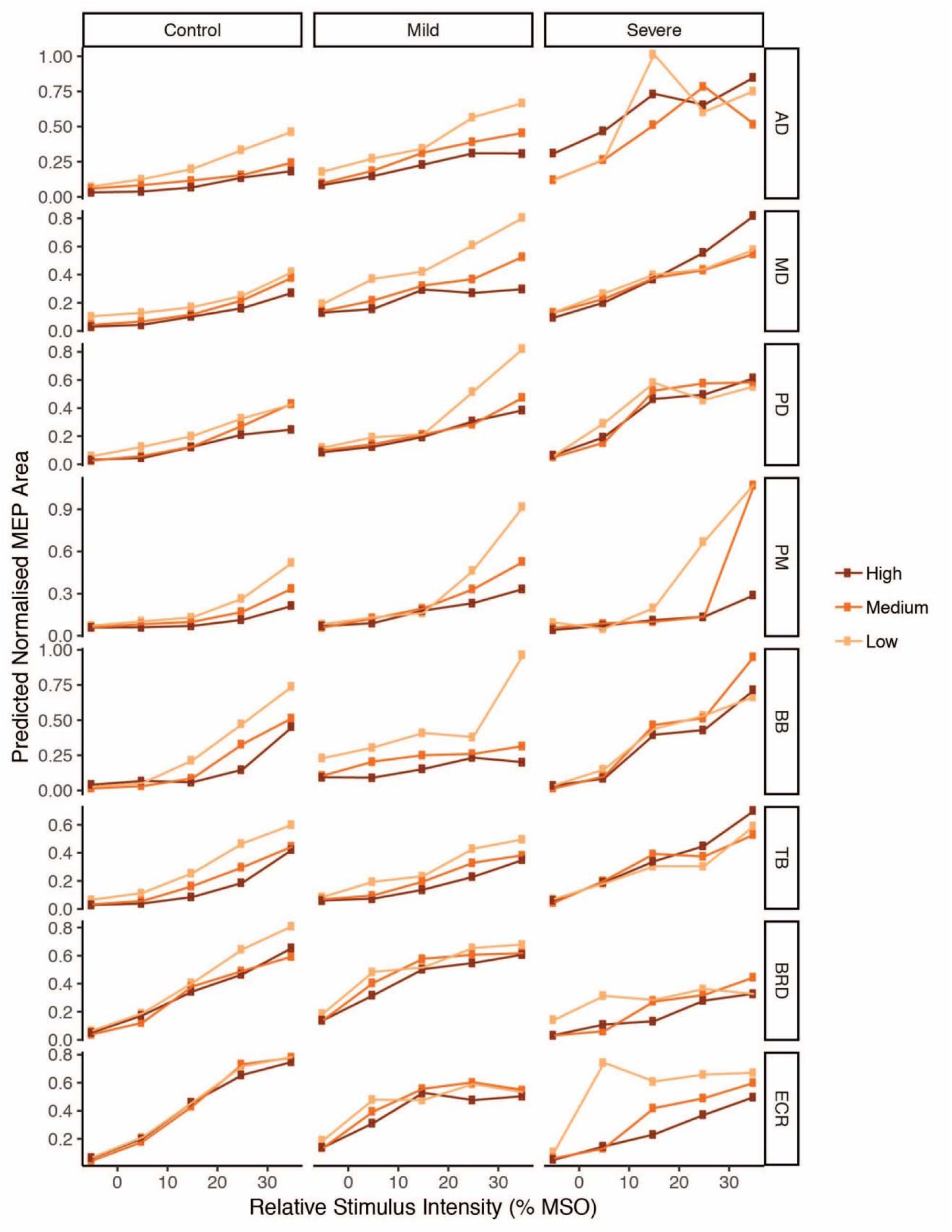
Predicted mean normalized MEP area is plotted as a function of stimulus intensity at low, medium, and high levels of support for control, mild, and moderate-severe impairment groups. Line shading represents support level

## Discussion

In this study we examined the interaction between upper limb impairment severity and WS during a reaching task. In support of our hypothesis, there was an interaction between impairment severity and WS on the number of targets hit. As expected, WS had the greatest effect for the moderate-severe impairment group who successfully reached an average of 2.5 more targets with high compared to low WS. An average difference of 0.1 targets for the mild impairment group and 0 targets for the control group is indicative that most participants in these groups could access the entire workspace without assistance. In line with previous findings, there was a significant effect of WS on iEMG for all muscles except TB (Prange et al., 2009a; 2009b; Coscia et al., 2014). Those with more severe impairment tended to exhibit smaller magnitudes of iEMG change between support levels. As expected, a muscle synergy analysis revealed that patients with worse impairment exhibited fewer synergies than healthy controls. Individuals in the control and mild impairment groups expressed more synergies with WS, indicating that WS facilitated more complex motor control (Figure 5). The number of synergies expressed by the moderate-severe impairment group did not respond to changes in WS, likely because of neuroanatomical constraints on available substrates for motor control. In the control group, WS affected CME (MEP area) measured in all muscles except ECR, similar to previous studies with WS and younger participants (Runnalls et al., 2014; 2017). Although WS influenced CME in both impairment groups, there was not a consistent pattern across muscles (Table 4, Figure 9). Taken together, these findings provide evidence that WS can influence the upper limb at behavioural and neurophysiological levels across the spectrum of motor impairments resulting from stroke.

### Direct and indirect effects of weight support

The direct mechanical action of WS is to reduce the magnitude of anti-gravity torques required at the shoulder. For the AD, MD, PD, and PM muscles, WS significantly lessened both iEMG during reaching (Supplementary Table 1) and rmsEMG during static abduction (Table 3). As expected, high targets required more activity than low targets, and far targets required more activity than near targets. This pattern was evident across levels of impairment (Figure 4). Impairment severity interacted with WS during the static abduction task; the moderate-severe group exhibited less modulation of rmsEMG between support levels (Figure 8). Similarly, the moderate-severe group exhibited less iEMG only with high support. In contrast, the control and mild groups tended to exhibit a more linear response between iEMG and support level similar to previous reports of healthy adults (Coscia et al., 2014). Impairment-dependent responses to WS could result from the recruitment of different neural elements for control. CME was also modulated linearly by WS only in the control and mild groups. Muscle activity patterns for the moderate-severe group are not reflected in the CME data. This disconnect could be a consequence of an increased reliance on alternative motor pathways to drive the proximal upper limb in individuals with significant corticospinal tract damage (Turton et al., 1996; Bradnam et al., 2012). The modulation of muscle activity in AD, MD, PD, and PM is primarily related to the direct mechanical effect of WS on shoulder joint torques. In control and mild groups, the up-regulation of CME with less WS appears to subserve voluntary drive to the proximal upper limb. In the moderate-severe group, neural drive may be distributed through alternative descending pathways that do not necessarily reflect modulation with WS as change in CME.

Dissociation of elbow muscle activation patterns between dynamic and static tasks provides evidence to support a distinction between direct and indirect effects of WS. A direct mechanical effect of WS is evident for the dynamic reaching task, where the elbow flexors BB and BRD acted against gravity and were sensitive to changes in WS. In contrast, TB was not oriented to act against gravity, thus compensating for gravity with WS was unlikely to have an effect on its activation. This was borne out by the absence of a WS effect on TB activity, which is consistent with previous studies of healthy adults (Prange et al., 2009b; Coscia et al., 2014), and stroke patients (Prange et al., 2009a). Elbow extension for reaching movements may be passively driven by the weight of distal segments, relying on modulation of antagonist elbow flexor tension to permit lengthening. An impaired ability to appropriately modulate BB iEMG for elbow extension may be reflected in the interaction of WS with impairment severity and target height. As expected, high targets required significantly greater BB activity; however, WS affected the groups differently. The application of medium support was sufficient to achieve maximum reduction of BB activity for the control group whereas the moderate-severe group required high support to achieve any change. In both cases, the application of WS through a forearm brace reduced the required force and had the expected effect of directly lessening BB activity.

In contrast to the direct effect of WS on BB activity observed during the reaching task, changes in BB and TB activity with WS during the static arm abduction task reflect an indirect effect of WS (Table 3, Figure 8). Whereas WS directly impacted the force required for shoulder abduction, BB and TB were oriented to act orthogonal to gravity and were not required to actively perform the task. However, the observed persistence of tonic activity in both muscles and its modulation with WS suggest a distinct mechanism may be responsible. An indirect effect of WS is further supported by changes in CME. The observed modulation of CME with WS in the BB and TB of healthy adults is consistent with previous experiments (Runnalls et al., 2014; 2017). Considering again the task did not require any activity at the elbow, the up-regulation of CME with less WS is likely an indirect effect of altered voluntary drive to proximal muscles mediated through neural linkages. Given that modulation of CME to distal muscles with WS depends on severity of impairment (Table 4), intracortical networks susceptible to disruption by stroke are implicated as a substrate for indirect WS effects. Whether the regulation of CME reflects a functional network architecture or incidental latent connectivity is unknown. In cases of less severe corticospinal system damage, ipsilesional motor cortical areas may provide a substrate well suited for neural reorganization subserving recovery. More severe impairments are associated with the recruitment of remote or secondary motor areas (Cramer et al., 1997; Johansen-Berg et al., 2002; Frost et al., 2003; Fridman et al., 2004). The resulting networks may be less efficient at generating motor output (Ward et al., 2006; Grefkes and Fink, 2011), or involve up-regulation of latent ipsilateral motor pathways with more diverse patterns of innervation (Bradnam et al., 2012). Mechanism aside, an indirect effect of WS was seen in the modulation of elbow muscle activity during a task in which they were not mechanically involved in gravity compensation and this modulation was smaller with greater severity of impairment.

Variation of muscle activity and CME in distal muscles that are not mechanically involved in gravity compensation provides evidence for an indirect effect of WS. In this study, the orientation and role of ECR was constant for both the reaching and static abduction tasks. However, manipulation of WS altered the amounts of iEMG during reaching and rmsEMG during static abduction (Tables 2 & 3). Similarly, greater iEMG was observed for both high targets and far targets despite requirements for wrist extension not varying between target locations. Modulation of muscle activity in ECR, which was mechanically unrelated to the action of WS, mirrored patterns observed in the proximal upper limb. The present findings suggest WS may influence distal muscle activity indirectly because distal muscles like ECR are subject to common neural drive carried over from proximal muscles.

The interaction of impairment severity and WS for ECR activity and CME was unexpected. During the reaching task, iEMG in the moderate-severe impairment group was lessened only with the highest level of support. Medium support was sufficient to achieve a similar reduction of iEMG for mild impairment. Overall, the patient response may be indicative of a WS threshold phenomenon. The control group paradoxically exhibited the most ECR activity with WS. The cause of this is unclear. It is possible the intact motor system has sufficient physiological range to permit some variation of activity in muscles that do not impact task outcome. A similar argument could explain why ECR CME did not respond to WS in the control and mild groups (Table 4). In contrast, previous experiments with healthy adults found that ECR CME was modulated by WS (Runnalls et al., 2014; 2017). Although it is unknown which factors may account for the discrepancy between studies, the present findings provide evidence in support of an indirect modulation of neural excitability distinct from changes in muscle activity.

### Impairment severity and mechanisms for integrated upper limb control

Integrated control of the upper limb based on neural linkages or synergies may facilitate the coordination of voluntary actions like forward reaching. In the present study, indirect effects of WS and interactions with impairment severity provide evidence for integrated control along the proximal-distal axis. Previous reports of distal CME modulation with changes to shoulder activation or shoulder position have interpreted the findings as task-relevant priming for muscle activation (Devanne et al., 2002; Dominici et al., 2005; Ginanneschi et al., 2005; 2006). The present findings support a model in which voluntary drive to proximal muscles acts as a regulatory signal in a proximal-distal hierarchy. Instances of dissociation between CME modulation and muscle activity (cf. Figures 6 & 7) suggest multiple neural linking mechanisms may be involved (see Runnalls et al., 2015). Along the same lines, different patterns of modulation between muscles could reflect the existence of multiple synergies with complex or competing behaviours. Differential responses depending on impairment severity provide a further indication that integration of control may be accomplished at many levels of the neuraxis.

Indirect responses to upper limb WS after stroke will depend on the neural structures disrupted by the stroke and whether the lesion is up- or downstream of the point where muscle activation information is linked together. Cortical binding of motor commands may be mediated in primary motor cortex where anatomical comingling of muscle representations may facilitate functional interaction (Sanes et al., 1995; Devanne et al., 2006). Proximal influences on distal CME may involve both intracortical disinhibition (Devanne et al., 2002; Kantak et al., 2013) and intracortical facilitation (Ginanneschi et al., 2005; 2006). Subcortical binding of motor commands may be mediated by divergence of descending corticomotor pathways (McKiernan et al., 1998), recruitment of the cortico-reticulo-propriospinal pathway (Pauvert et al., 1998; Pierrot-Deseilligny, 2002), or activation of spinal interneuron modules (Bizzi and Cheung, 2013). Subcortical lesions at the level of the PLIC would disrupt the pattern of cortically linked neural activity and the activation amplitude of subcortically linked neural activity (McMorland et al., 2015). Any combination of these factors may affect the coordination of descending neural information.

### Effects of weight support on muscle synergy expression

Control and mildly impaired participants were able to recruit more complex patterns of muscle activity with WS. The emergence of a higher number of synergy clusters with greater support suggests that WS facilitates independent muscle recruitment. The emergence of new synergies should allow for greater functionality and thus potentially benefit rehabilitation therapy. In contrast, the moderate-severe impairment group expressed constant muscle synergies at all levels of support. Although WS increased the number of targets hit by this group, it did so without altering the underlying structure of muscle activity. These results, considering the smaller task space, support the idea that more severe damage after stroke leads to a reduced number of synergies (Clark et al. 2010). The number of muscle synergies therefore provide a possible marker of neuroanatomical damage (Cheung et al. 2012).

Synergy structure was conserved across levels of support even in the presence of neural damage (Israely et al. 2018). Furthermore, synergies in patients with mild impairment were similar to controls (Cheung et al. 2009), exhibiting stereotypical changes previously described (Roh et al. 2015). Three synergies were identified for the control group: external rotation, internal rotation, and flexion synergy. The use of these three synergies aligns well with the task space. The task space constrains the number of synergies and may explain why fewer synergies overall were seen compared with other studies (Roh et al. 2012). Two synergies were identified for the mildly impaired group: an atypical external rotation compared with the control group, and internal rotation. Finally, just one atypical synergy was identified for those with moderate-severe impairment. It is noteworthy that for the moderate-severe group, WS increased the number of targets reached and modulated the magnitude of muscle activity without affecting the number of synergies expressed. This observation demonstrates a dissociation between the amount of muscle activity and the structure of that muscle activity. This could reflect more severe damage to neuroanatomical substrates constraining the neuromechanical repertoire. In contrast, the control and mild groups expressed more synergies with WS while at a task performance ceiling. This latter observation could indicate that WS can facilitate access to latent regions of the motor control space and improve movement quality by enabling more efficient muscle activity. Whether such phenomena can be meaningfully exploited for neurorehabilitation will be an interesting avenue of future research.

### Potential limitations

A limitation of the present study is the absence of kinematic measures of reaching performance. A quantitative characterization of movement quality could reveal additional effects of WS and add context to the interpretation of EMG data. The reaching task, as defined by the array of targets, was designed to accommodate individuals with a broad range of impairments. However, there was a trade-off in terms of sensitivity to detect changes. Future studies may wish to incorporate more gradations in target location or additional constraints such as a retrieval component. The present study used procedures suitable to obtain only contralateral MEPs from stimulation of the ipsilesional hemisphere. Additional measures, e.g. ipsilateral MEPs, may have yielded neurophysiological data from more of the patients with severe CST damage.

## Conclusions

Arm weight support may benefit stroke patients with upper limb impairment through both direct and indirect neural mechanisms. First, by directly lessening forces required to complete tasks, individuals with decreased force generating capacity can access a larger workspace and engage in practice with a wider range of tasks. Second, by indirectly influencing linked neural elements, arm weight support may promote a rebalancing of corticomotor excitability in otherwise saturated networks. Potentially, individuals can then engage a neurophysiological landscape more permissive to modulation and plasticity. The threshold of weight support required to achieve a desired modulation will vary between muscles and tasks, and almost certainly depend on the extent of impairment. Keeping these factors in mind, weight support may be a useful adjuvant to upper limb rehabilitation after stroke.

## Acknowledgments

The authors acknowledge assistance provided by April Ren, Terry Corin, Fiona Doolan, and support from Saebo Inc. for supplying the SaeboMAS. WB received funding from the Health Research Council of New Zealand.

## Supplementary Material

**Supplementary Table 1.**
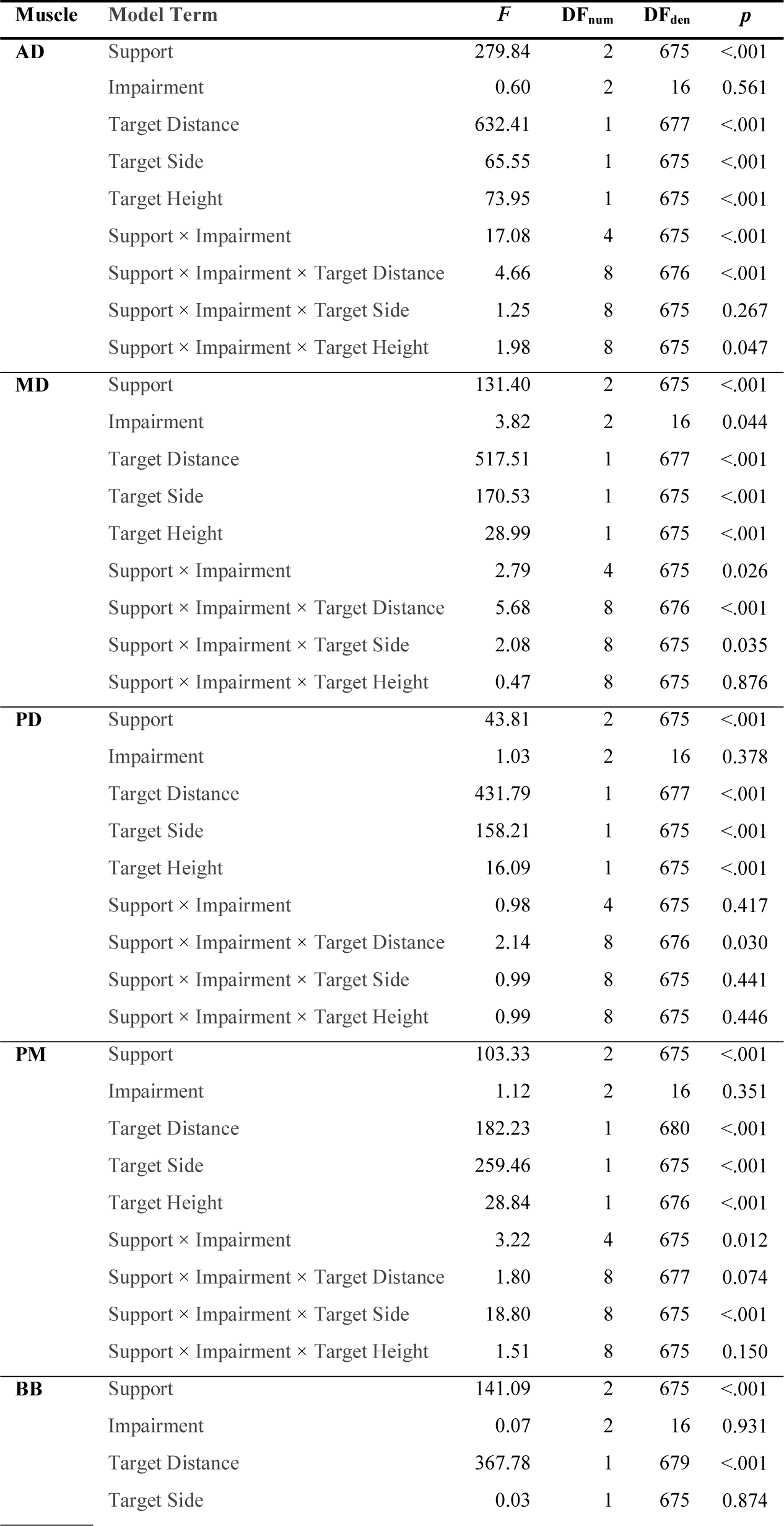
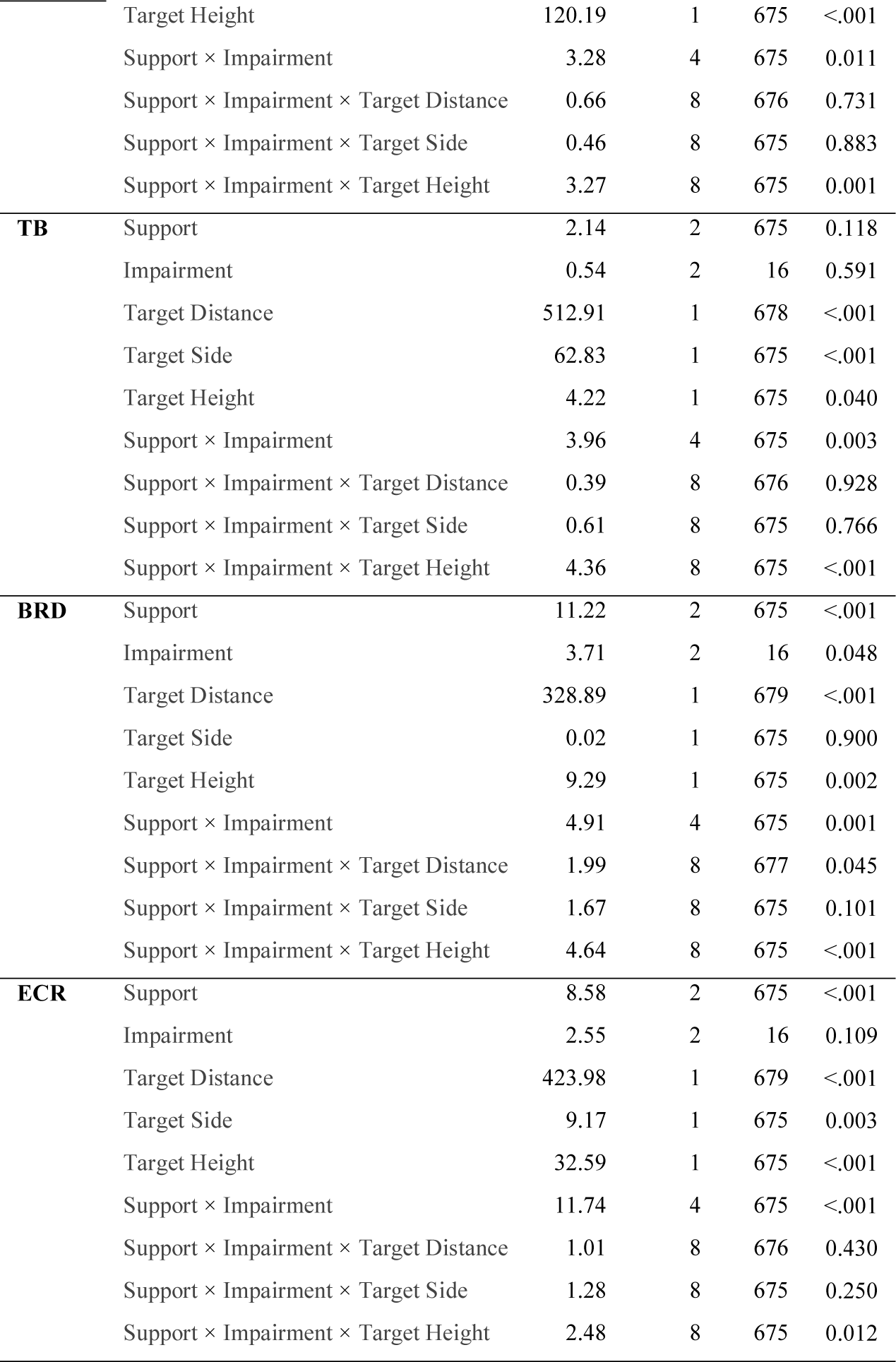
ANOVA for linear mixed models of iEMG during reaching task

